# Antibody mediated epitope mimicry in the pathogenesis of Zika virus related disease

**DOI:** 10.1101/044834

**Authors:** E. Jane Homan, Robert W. Malone, Steven J. Darnell, Robert D Bremel

## Abstract

The association of Guillain-Barré syndrome with Zika virus infection raises suspicion of autoimmunity in the pathogenesis of Zika associated disease. Using computational analysis to identify predicted B and T cell epitopes, we assessed whether antibodies elicited by B cell epitopes in Zika virus may also target B cell epitopes in the human proteome. We detected amino acid motifs predicted to be B cell epitopes in Zika virus proteins which are also present in human proteins, including pro-neuropeptide Y (proNPY), NAV2 and other proteins with interacting neurophysiologic function. We examine the predicted MHC binding of peptides likely to provide T cell help to the potential mimic epitopes. Some potential mimic epitopes in Zika virus envelope have apparently strong T cell help, likely facilitating immunoglobulin class switch. We also identify epitope mimic commonalities with dengue serotypes 1 and 3. We hypothesize that antibodies to Zika virus epitopes may contribute to the pathogenesis of Zika-associated Guillain-Barré syndrome, microcephaly, and ocular lesions, and may be a driver of autoimmunity. The risk associated with responses to potential epitope mimics must be addressed in the development of vaccines and therapeutics for Zika virus infections.

**Author Summary:** Using computational immunologic analysis, we examine the possibility that anti-Zika antibodies are binding to mimic epitopes on human proteins and that this autoimmunity may be a driver for some of the clinical signs associated with Zika virus infection. These include Guillain Barré Syndrome, other neurophysiologic deficits, and the Zika Fetal Syndrome, including microcephaly. We identify specific proteins and epitopes to which anti-Zika antibodies may bind. The prospect that the pathogenesis of ZIKV may involve an antibody-mediated autoimmune component must be addressed in vaccine and therapeutic development.

## Introduction

Zika virus (ZIKV) has emerged as a rapidly spreading epidemic throughout the tropical Americas. Known to cause mild febrile disease since its discovery in Uganda almost 70 years ago [1], changes in the clinical signs of Zika were noted following its appearance in French Polynesia, and arrival in Latin America. Here, infection may be followed by Guillian-Barré Syndrome (GBS)[2] and adult infection is also associated with other sensory neurologic impairment [3]. Spontaneous abortion and primary microcephaly, following infection during the first 2 trimesters of pregnancy, is increasingly regarded as caused by ZIKV infection[4, 5]. Ocular lesions have been noted in surviving infants [6, 7]. These and other neonatal neurologic sequelae of infection during pregnancy constitute Zika fetal syndrome. GBS has been reported sporadically following dengue infection [8, 9], but is reportedly more severe in Zika case [10]. The incidence of teratogenic lesions sets Zika apart from dengue. ZIKV has been detected by PCR in fetuses, and high concentrations of Zika virion particles have been detected by electron microscopy in the fetal central nervous system [4]. The combination of GBS and Zika fetal syndrome associated with infection suggest that these disease syndromes may share common etiology such as an autoimmune response targeting neurologic functions, and raises the possibility of epitope mimicry [11].

Seeking to illuminate Zika pathogenesis, in the studies described herein we have focused on potential autoimmune pathways rather than considering the virus as the only agonist of pathology. We have utilized an applied mathematical approach to evaluate predicted immune responses to Zika. We compare the profile of predicted antibody mediated epitope mimicry of ZIKV in the Americas with that of Old World ZIKV. We have predicted proteins in the human proteome, with functions in neurophysiology and fetal development, and which may be functionally compromised by binding to anti-Zika antibodies.

## Methods

### Virus sequences

Polyprotein sequences were downloaded from Genbank and manually curated into gene product proteins based on GenPept annotations. To assess how the pathogenesis of ZIKV related disease changed between Africa and the Americas, our effort focused initially on comparing two representative isolates: Brazilian isolate SPH2015 (gi: 969945757, “Zika Brazil”) [12] and a ZIKV isolate representative of the West African clade (gi: 592746966 “Zika Africa”) [13]. Subsequent analyses were conducted on other ZIKV isolates, and on related flaviviruses, including recent South American isolates and reference strains of dengue, yellow fever, and representative neurotropic flaviviruses. Flaviviruses analyzed are listed in Supporting Table 1.

Given the co-endemnicity of chikungunya virus, and because of similarity of the microcephalitic lesions linked to Zika to those following intrauterine exposure to cytomegalovirus (CMV) and rubella, we also performed a preliminary analysis of structural proteins of a small number of these viruses. Sequences for chukungunya and rubella were accessed from NIAID Virus Pathogen Database and Analysis Resource (ViPR) [14], using 10 geographically distinct but random wild type isolates of each. In addition, a complete proteome of CMV Towne was analyzed (FJ616285).

### Proteome

The human proteome dataset was downloaded from the UniProt repository (Nov 2013) [15]. It was manually curated to remove sequences derived from antibody V-regions. UniProt continually re-curates and updates the repository, but changes in the database since downloading are inconsequential for the use described herein. As curated in our laboratory, the dataset represents slightly greater than four-fold redundancy and contains a total of 88,145 identified proteins comprising 3.498 x 10^7^ peptides.

### Epitope analysis

Binding of viral peptides to B and T cells is a competitive process. To assess binding in the competitive context of each protein, the polyprotein of each flavivirus was broken out into individual proteins based on GenPept annotations to allow standardization of competitive binding. Arrays of predicted major histocompatibility complex (MHC) binding, as well as of linear B cell epitope probability were constructed for each sequential peptide in each of the proteins. The applied mathematical tools used for the immunoinformatic analysis have been previously described [16, 17]. Briefly, ensembles of neural network (NN) prediction equations were developed using publically available databases of MHC class I and class II binding data [18, 19] using the neural platform of the JMP^®^ (SAS Institute, Cary, NC) statistical software application. As initially described by Wold *et al* [20], for multiple regression predictions the first three principal components of amino acid physical properties were used as predictors in the input layer of the neural networks. Ten ensembles are used for each allelic prediction. The use of ensembles enables estimation of the mean binding affinity and an estimate of the variation in the NN binding affinity prediction. All binding data generated by the NN were transformed to a within-protein zero mean unit variance distribution using a Johnson Sb distribution algorithm in JMP^®^. This transformation puts all binding data on a common scale, making it possible to use the principle of additivity of variance.

### B cell epitope prediction

Viral proteins were analyzed to compute predicted probability of B cell epitopes (BEPIs), standardized within protein. We had precomputed the proteome database to determine peptides that fulfil the criteria of a potential BEPI within each source protein. We use a sliding 9-mer window to compute predicted BEPIs, and for comparative purposes, the pentamer central core of each predicted BEPIs was used to seek matches. A pentamer is chosen because, not only does it provide a very stringent filter, but it corresponds to the area needed to engage the paratope of an antibody [21]. A series of filters was applied to (a) find ZIKV protein pentamers with the highest probability of functioning as BEPIs, this was initially set at the top 25%, and (b) identify pentamers with the highest probability of being BEPIs in each of the >88,000 human proteins, initially set at the top 40%. Matching BEPI pentamers were identified, then a third filter applied (c) a search term of key words to extract human proteome proteins with neurologic functions. In this case we utilized keywords comprising 56 variations on the terms “neur”, “glial”, “myelin”, “opt”, and “synapt” (Supporting Table S2).

Pentamers fulfilling all of these three criteria were declared to be potential Zika-proteome epitope mimics. The stringency of these criteria can then be increased to identify the highest probability mimics. This process provides a highly selective set of filters. Any single pentamer has a 1 in 20^5^ chance of occurrence (5 of 20 amino acids). When this probability is applied independently to both the constituents of the Zika polyprotein and to the human proteome sets, there is a 3423/20^5^x20^5^ chance of a match, or 1 in 3.3x10^10^. This probability is further reduced by application of the BEPI and keyword filters. However, the occurrence of multiple isoforms of some proteins increases probability. In a further independent evaluation of the ZIKV proteins, the adjacency to probable BEPIs of 15-mers with predicted high affinity MHC II binding was determined, as these may stimulate specific T cell help. T cell help may stimulate a higher titer of antibody, and will enable class switch to IgG.

The characteristics and performance of our B cell epitope (BEPI) prediction tools have likewise been described elsewhere [22]. Briefly, like the MHC predictions, the BEPI prediction tools comprise NN and use amino acid physical property principal components, and are aligned to the BEPIPred [23] approach,. and developed to work within our data processing workflow. The essential prediction is a probability that the central amino acid in a nonamer peptide resides within a combinatorial “patch” of amino acids that might be a BEPI. The probability is transformed to a within-protein zero mean unit variance distribution using a Johnson Sb distribution. This process puts the metrics onto a common scale making it possible to use the scaled metric as a statistical screening criterion that can be used on groups of proteins. For convenience of comparison to standardized MHC binding where negative standardized scores represent high affinity binding, the standardized BEPI probabilities are inverted (multiplied by −1).

Whereas MHC binding predictions lead to a specific binding affinity prediction for a particular peptide, this cannot be done in the case of a B cell somatic hypermutational response in a germinal center response to infection. The B cells will generate an enormous number of different antibody molecules with differing affinities targeting the same “patch” of amino acids. Moreover, as has been shown for the B cell repertoire responders to tetanus toxoid the population of B cells surviving in the steady state may represent a small fraction of the early responding population [24]. We have restricted the primary screen to the central pentamer of the nonamer in the NN. A central pentamer can be found in different molecular contexts with different predicted propensities for B cell interactions. Such a central pentamer can be found in 20^4^ different flanking tetramer contexts with different predicted propensities for B cell interactions. The Pearson correlation (r = 0.44, P < 0.0001) measures the magnitude of the effect of the flanking tetramers on overall BEPI probability (Supporting Figure S1).

During the analysis it became apparent that ZIKV pentamers with high BEPI probability were also generating very high probability matches to von Willebrand’s factor A (VWA). Because this factor could be implicated in the clinical signs (hemorrhagic microvascular rash), this group of pentamers was added to the subset for further analysis.

Thus, we predicted ZIKV BEPIs likely to give rise to anti-Zika antibodies, and to also specifically react with BEPIs in target human proteins of neurologic or other selected functions. This process was repeated with other ZIKV isolates, and the other selected flaviviruses. The predicted Zika-proteome mimics identified by the screening process were evaluated to address the following questions: How do epitope mimics in Zika Brazil differ from those predicted in Zika Africa? Are potential mimics unique to Zika versus other flaviviruses? Do other flaviviruses have mimics in the same human proteins? What fraction of the MHC alleles are likely to provide help in generating antibodies targeting these mimics?

### Structural Analysis

Envelope (E) protein structure models were predicted using NovaFold® (DNASTAR, Inc., version 13.0, structure template library 2015-W47-5). NovaFold®, which is based on the I-TASSER structure prediction algorithm [25-27], selects a collection of potential structure templates using protein threading alignments, then perturbs the selected templates into lower-energy conformations using replica exchange simulations, and finally relaxes the backbone and side chains of the final models using all-atom energetic refinement. The E protein ectodomain sequences for Zika Brazil (GenBank: ALU33341.1) and Zika Africa (GenBank: AHL43504.1) strains were obtained from the Zika virus polyprotein (residues 291-690). The top templates included E protein structures from West Nile (PDB ID: 2I69) and Tick-borne encephalitis viruses (PDB ID: 1SVB). Accuracy is estimated in terms of predicted Template Model score

(TM-score; a metric weighing close atom pairs between structures greater than more distant atom pairs; closer to 1 is better, greater than 0.5 indicates a correct fold) [28, 29]. TM-score has been proven to be a more robust metric than root mean square deviation (RMSD) as it is less sensitive to local structural variations [30]. For the Brazil model, the predicted TM-score was 0.85; for the Africa model, it was 0.87. Both scores are indicative of a high degree of confidence in the accuracy of the predicted structures.

Surface patches of the Zika virion were modeled by aligning individual copies of the predicted E protein models onto the cryo-electron microscopy structure of the dengue serotype 1 virus (PDB ID: 4CCT, resolution 4.5 Å). The modeled virion patch consists of the following dengue E proteins from the biological assembly: chains A, B, and C from model 1; chains A and C from model 6; and chain B from model 10. Pairwise structure alignments were performed using TM-align [31] in Protean 3D™ (DNASTAR, Inc., version 13.0). Compared to the dengue E protein structure, the TM-score was 0.77 for the Zika Brazil E protein model and 0.74 for the Africa model. Solvent-accessible surfaces for the collective patch of six E proteins were calculated using EDTsurf [32] in Protean 3D™.

## Results

### Envelope protein epitopes

As the structural proteins are exposed to the immune system and account for nearly all of the antibodies elicited by comparable flaviviruses, we first examined the structural proteins, with emphasis on the envelope protein [33, 34]. Despite substantial sequence differences in the envelope proteins among the different viruses, there is a strong similarity in the molecular location of predicted linear B cell epitopes (Figure 1). Our in silico predictions of dengue linear B cell epitopes are consistent with those mapped by monoclonal antibodies [35, 36]. Epitopes in envelope domain II (EDII), conserved across other flaviviruses and thought to contribute to antibody dependent enhancement (ADE) [37], are also preserved in Zika virus. A band of predicted high affinity MHC binding, located at amino acid positions 194-224 (Figure 1), is observed in all flavivirus envelopes examined. However, there is more difference in distribution of other peptides with predicted high affinity MHC class I and II binding between the analyzed flaviviruses. Of note in ZIKV is a region at amino acid positions 369-390, adjacent to the DE loop of envelope domain III (EDIII), with high predicted MHC II binding across all HLA alleles.

**Figure 1.**
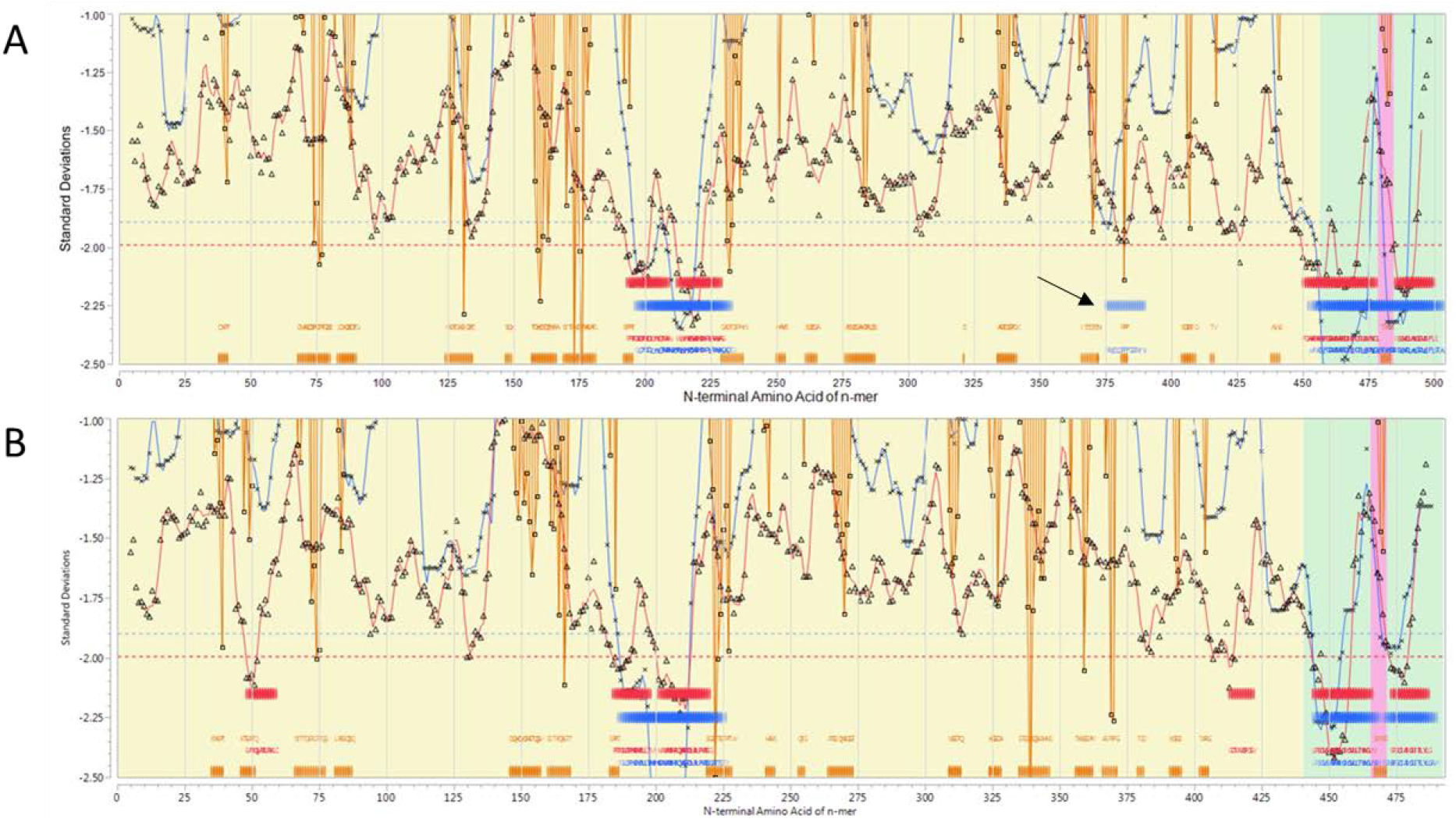
Comparative linear immunome of the envelope proteins of Zika Brazil and dengue 3. A. Zika Brazilian isolate SPH2015 (gi: 969945757); B. Dengue 3, a 2009 Brazilian isolate D3BR/AL95/2009 gi: 389565793. X axis shows N>C amino acid positions. Y Axis shows standard deviation units of predicted MHC binding. Background shading shows transmembrane domains (green) outside envelope (yellow), intraenvelope (pink). Predicted MHC-I (red line), MHC-II (blue line) binding, and probability of B cell binding (orange lines) for each peptide, arrayed N-C, for a permuted population comprising 63 HLAs. Ribbons (Red=MHC-I, Blue-MHC-II) indicate the top 25% affinity binding. Orange bars indicate high probability B-cell binding. Arrow indicates band of predicted MHC II binding in the Domain 3 DE loop of Zika envelope.

Overall, the concordance between our in silico predictions and traditional experimental epitope mapping in flaviviruses lends support to the approaches described below, and is consistent with prior reports [22]. We also noted (data not shown) that compared to other viruses of similar size, ZIKV has an overall very high degree of peptide motif match to the human proteome.

### Predicted epitope mimics in Zika Brazil vs Zika Africa

Conformational B cell epitopes undoubtedly play an important role in ZIKV immunology, and may cross link molecules and domains of the envelope protein, as with dengue [34]. However, the simplest form of molecular mimicry would be an exact match of a group of amino acids similarly exposed as a linear B cell epitope in both a viral and a human protein. In such cases, viral infection may stimulate antibodies that also bind B cell epitopes in the human proteome. Collectively, all ZIKV proteins in the two isolates examined comprise 103 pentamers meeting the criteria of predicted Zika-proteome epitope mimics laid out above. Of these, 88 were conserved between Zika Africa and Zika Brazil, 15 distinct new ones were gained in Zika Brazil, and 13 present in Zika Africa were lost before arrival in Brazil. Table 1 summarizes those in the three structural proteins. Mimic matches for all proteins are listed in Supporting Table S3. Figure 2 shows five potential mimics stand out as having the highest probability of generating a high antibody titer.

**Figure 2.**
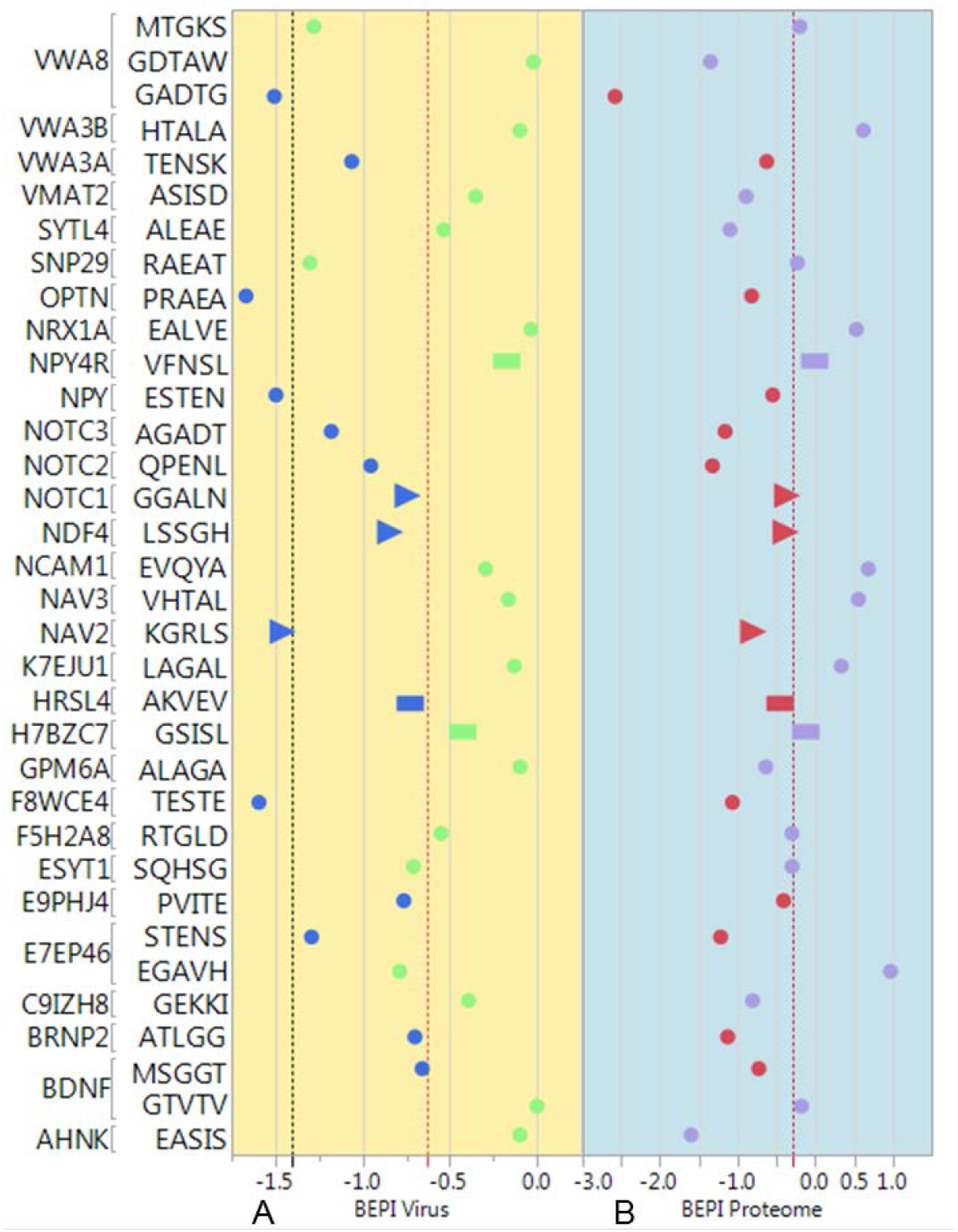
Categorization of potential mimic pentamers in the envelope proteins of Zika virus. Pentamers occurring in envelopes of the two Zika isolates examined are listed on the Y axis and grouped by human proteome neurologic protein match. X axis shows the probability of a pentamer being a B cell epitope (BEPI) in negative standard deviation units. Panel A shows predicted BEPIs in the virus envelope. The red dotted line identifies the top 25%; black line the top 10% cutoff; Panel B shows predicted BEPIS in the human proteome neurologic proteins, with the red dotted line indicating the top 40%. Round markers are motifs conserved in Zika Brazil and Zika Africa; triangular markers are in Zika Brazil only; bars identify motifs only in Zika Africa. Dark blue and red markers identify motifs meeting all selection criteria; pale green and lilac markers fail on one or more criteria of selection.

**Table 1.** Pentamer predicted B cell epitope mimics in the structural proteins of Zika Brazil and Zika Africa.

| Pentamer | Zika AFR | Zika BR | BEPI Virus | BEPI Proteome | UniProt ID | Annotation |
| --- | --- | --- | --- | --- | --- | --- |
| Envelope |  |  |  |  |  |  |
| **PRAEA** | Y | Y | -1.67 | -0.84 | OPTN | Optineurin |
| **TESTE** | Y | Y | -1.59 | -1.07 | F8WCE4 | Synaptogyrin-1 |
| **GADTG** | Y | Y | -1.51 | -2.32 | VWA8 | von Willebrand factor A |
| **ESTEN** | Y | Y | -1.50 | -0.55 | NPY | Pro-neuropeptide Y |
| **KGRLS** | N | Y | -1.46 | -0.80 | NAV2 | Neuron navigator 2 |
| STENS | Y | Y | -1.29 | -1.22 | E7EP46 | Neurotrophin-4 |
| AGADT | Y | Y | -1.18 | -1.16 | NOTC3 | Neurogenic locus notch homolog protein 3 |
| TENSK | Y | Y | -1.06 | -0.63 | VWA3A | von Willebrand factor A |
| QPENL | Y | Y | -0.95 | -1.32 | NOTC2 | Neurogenic locus notch homolog protein 2 |
| LSSGH | N | Y | -0.84 | -0.38 | NDF4 | Neurogenic differentiation factor 4 |
| PVITE | Y | Y | -0.76 | -0.41 | E9PHJ4 | Neural cell adhesion molecule L1 |
| GGALN | N | Y | -0.74 | -0.37 | NOTC1 | Neurogenic locus notch homolog protein 1 |
| AKVEV | Y | N | -0.73 | -0.46 | HRSL4 | Retinoic acid receptor responder protein 3 |
| ATLGG | Y | Y | -0.70 | -1.13 | BRNP2 | BMP_retinoic acid-inducible neural-specific protein 2 |
| MSGGT | Y | Y | -0.66 | -0.52 | BDNF | Brain-derived neurotrophic factor |
| PrM |  |  |  |  |  |  |
| ARRSR | Y | Y | -1.65 | -0.95 | NEUL2 | Neuralized-like protein 2 |
| SDAGK | Y | N | -1.46 | -1.55 | E7EUC6 | Neuron navigator 3 |
| GSSTS | Y | Y | -1.27 | -1.95 | SYPL2 | Synaptophysin-like protein 2 |
| STRKL | Y | Y | -1.15 | -0.59 | A2A341 | Synaptonemal complex protein 2 |
| SHSTR | Y | Y | -1.02 | -0.63 | F5GZS7 | Neuregulin-2 |
| RSRRA | Y | Y | -0.99 | -0.93 | ARHG8 | Neuroepithelial cell-transforming gene 1 protein |
| Capsid |  |  |  |  |  |  |
| KKRRG | N | Y | -2.21 | -1.69 | H7BY68 | Putative neuroblastoma breakpoint family member 8 |
| RRGAD | Y | Y | -2.11 | -0.75 | NEUL4 | Neuralized-like protein 4 |
| EKKRR | N | Y | -2.05 | -1.55 | NPAS2 | Neuronal PAS domain-containing protein 2 |
| ERKRR | Y | N | -1.95 | -0.60 | NSMF | NMDA receptor synaptonuclear signaling and neuronal migration factor |
| SVGKK | Y | Y | -0.93 | -0.61 | ESYT3 | Extended synaptotagmin-3 |

To examine the locations of the novel motifs identified by the predicted mimic pentamers, we constructed an atomic (3D) model of the Zika Brazil and Zika Africa envelope protein (E protein) using NovaFold^®^ structure prediction software from DNASTAR, Inc. NovaFold^®^ uses a hybrid modeling approach that incorporates homologous structures and *ab initio* modeling to accurately predict a protein’s 3D structure from its amino acid sequence. As seen in Figure 3, the pentamers contained within the Zika Brazil PVITESTENSK sequence are expected to reside on the solvent-accessible DE loop in EDIII[33]. These include predicted mimic matches for pro-neuropeptide Y (proNPY) (ESTEN), synaptogyrin (TESTE), neurotrophin 4 (STENS), neural cell adhesion molecule (PVITE), and VWA (TENSK). Adjacent to this is a sequence comprising overlapping 15-mer peptides that, in aggregate have predicted high affinity MHC II binding for almost all mapped DRB, DP and DQ alleles (Figure 4). Peptides from this region are thus likely to provide context-specific T cell help through co-presentation by B cells [38], supporting development of high antibody titer and class switching.

**Figure 3.**
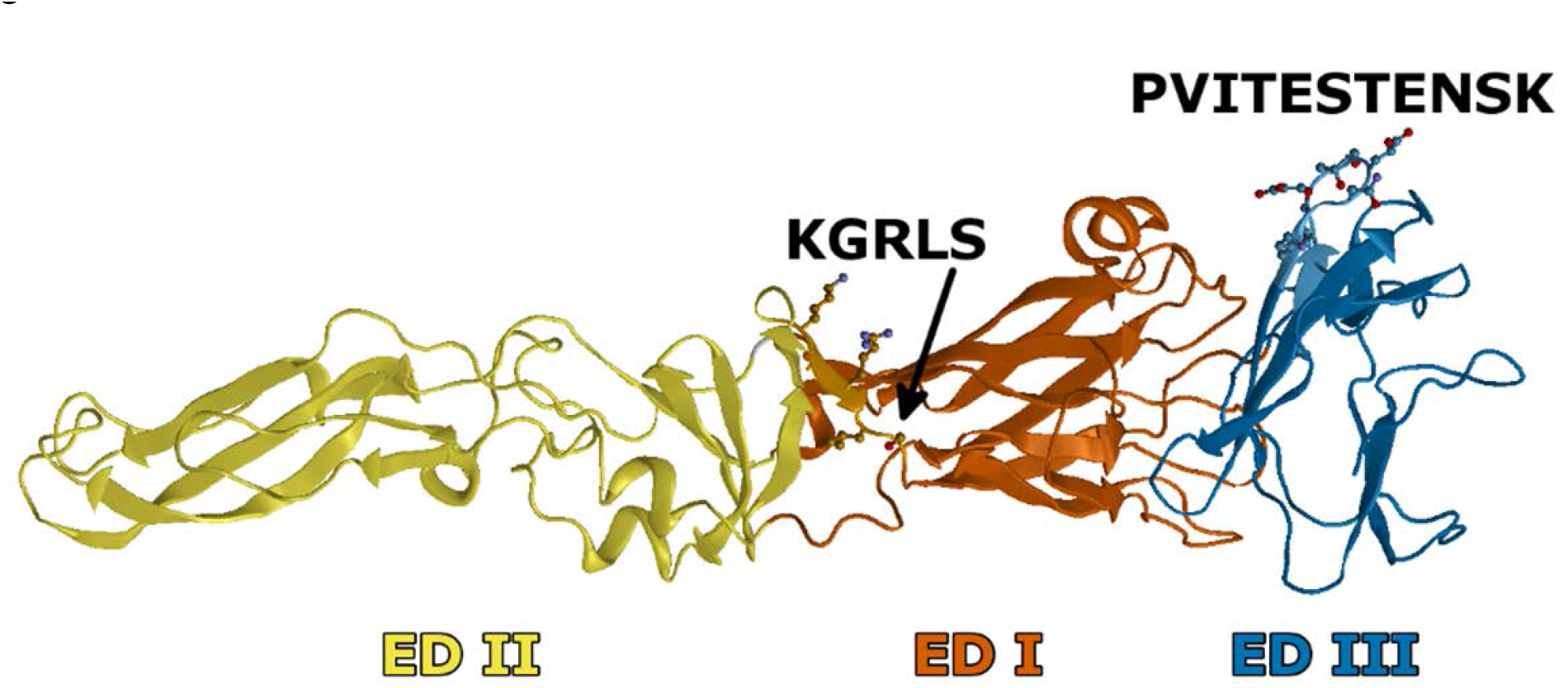
Model of the ectodomain for the Zika Brazil E protein predicted by NovaFold^®^ (DNASTAR, Inc.). The E protein domains (ED) are colored in red-orange (I), yellow (II), and blue (III). Side chains are displayed for residues associated with the labeled epitopes. The arrow for the KGRLS epitope indicates the position of Ser 285.

**Figure 4.**
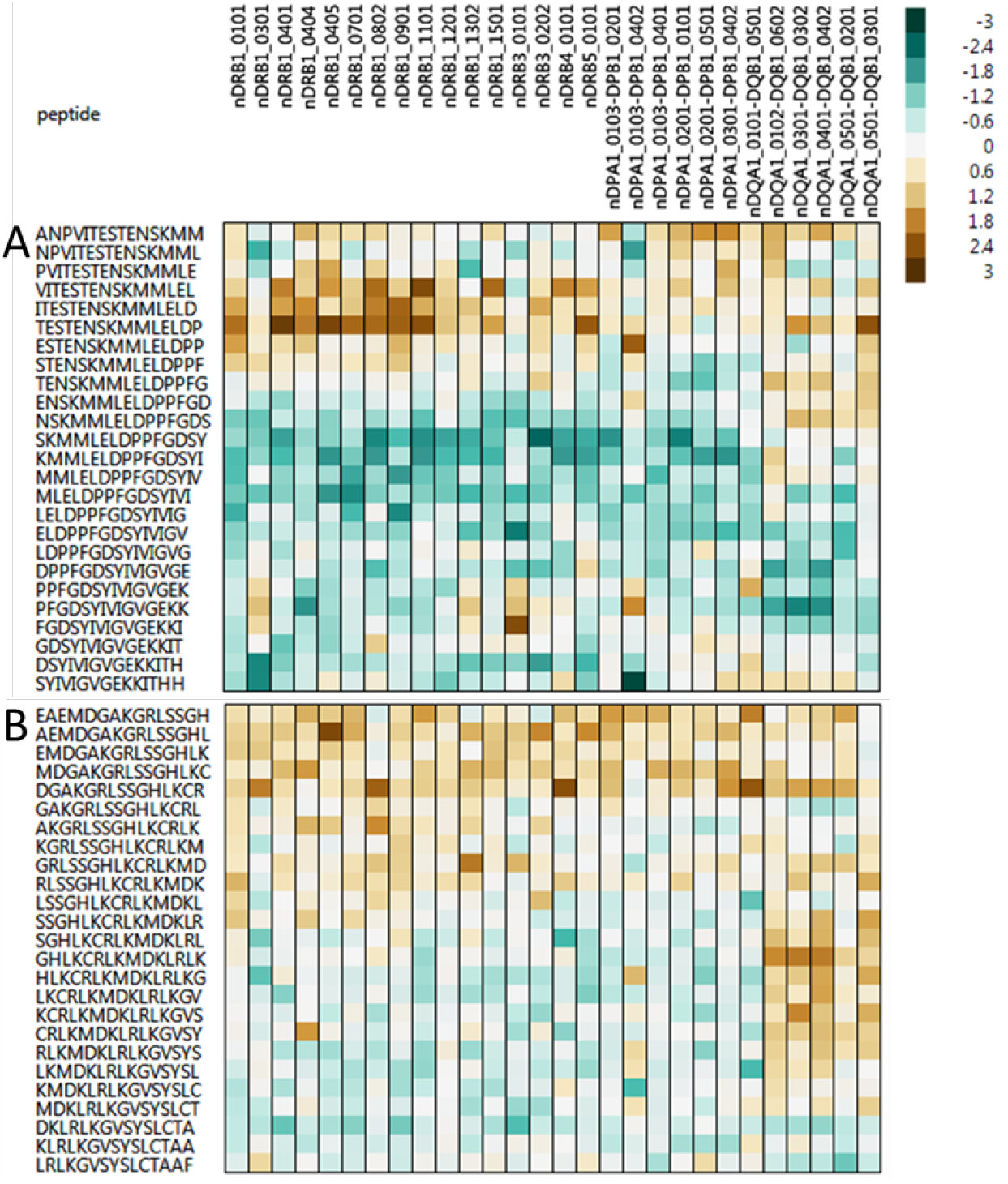
Predicted MHC II binding affinity adjacent to ESTEN and KGRLS X axis (top) shows HLA II analyzed. Y axis shows 15-mer peptides across the sequences of interest adjacent to PVITESTENSK (A) and KGRLS (B). Cell coloration shows predicted binding affinity in standard deviation units. Dark green is highest affinity predicted binding, brown lowest.

A high probability of a mimic to optineurin occurs in EDI (PRAEA). Structurally adjacent to this, EDI also comprises two overlapping predicted pentamer mimic motifs, found in Zika Brazil but not Zika Africa, with matches to neural navigator 2 (NAV2) (KGRLS) and to neural development factor 4 (LSSGH). These predicted new mimics arise from a single SNP leading to the F285S mutation, first detected in SE Asian isolates.

To further investigate the KGRLS epitope, we constructed a model of the Zika Brazil and Africa E proteins in a mature virion configuration (Figure 5) using Protean 3D™ molecular modeling software from DNASTAR, Inc. Both Ser 285 (Brazil) and Phe 285 (Africa) are expected to be solvent-accessible, but also reside in a cleft formed at the interface between two E protein subunits. The F285S mutation suggests this cleft may be deeper and more polar in Zika Brazil, altering the BEPIs exposed at this location. The virion models also suggest that, given their relative spatial proximity, Thr 156 and Ser 285 in Brazil and Ile 156 and Phe 285 in Africa may form conformational epitopes with differing geometric and chemical characteristics. A further predicted new mimic (GGALN), for neurogenic locus notch homolog protein 1 (NOTC1), occurs close to the C terminus of the soluble portion of the envelope, but is detected only at a low stringency BEPI probability selection.

**Figure 5.**
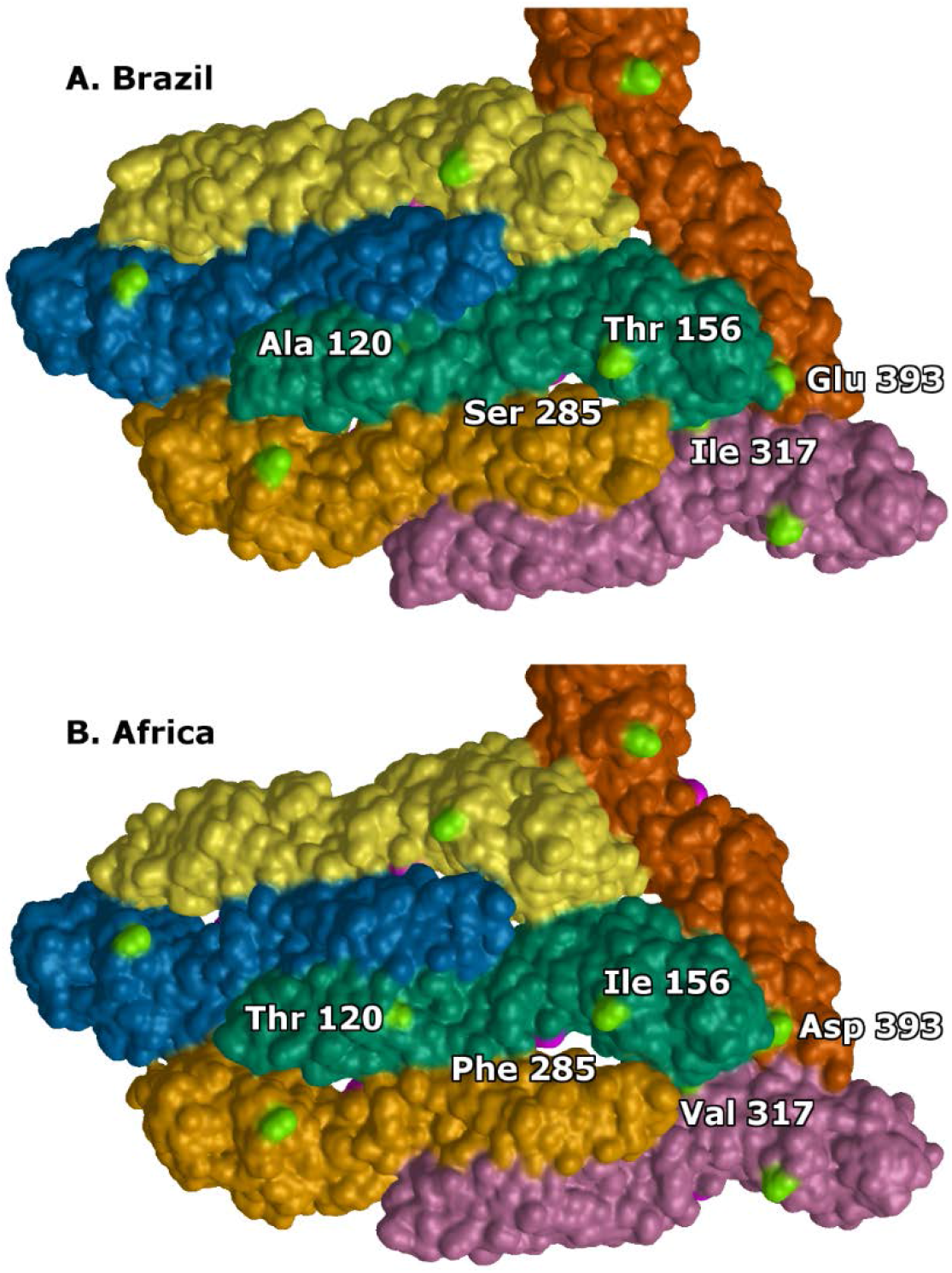
Model of a virion surface patch consisting of E proteins for Zika Brazil (A) and Zika Africa (B) variants. Predicted monomer models are structurally aligned to the biological assembly of the dengue serotype 1 virus structure (PDB ID: 4CCT) in Protean 3D™ (DNASTAR, Inc.). The solvent-accessible surface is colored to distinguish the location of the individual E proteins. Labeled residues on the central green subunit denote nonsynonymous variations between Zika Brazil and Zika Africa. Additionally, the surface is colored bright pink for region associated with position 285 (part of the KGRLS epitope) and light green for the remaining residues. Positions 156 and 285 are colored in the other subunits to establish the orientation relative to the central subunit. Notably, Ser 285 (A) is predicted to be solvent exposed, but residing in a cleft formed between two subunits.

### Epitope mimics in other flavivirus envelope proteins

We compared epitopes of ZIKV and other flaviviruses, using an approach designed to identify complex sharing patterns, to assess the differences in antibody mimics that each may generate. Despite the taxonomic similarity there are few predicted mimics in common between ZIKV and the other flaviviruses investigated. Only two of the pentamer predicted Zika mimics were found to be identical to mimics predicted from other flaviviruses. AGADT, shared by Zika Brazil, Zika Africa and dengue 4, does not fulfill the mimic criteria of BEPI probability in the proteome. QPENL, shared by Zika Brazil, Zika Africa and with dengue type 2, only does so at the lowest stringency.

The other flaviviruses do contain a small number of potential mimics, which could generate antibodies that target the same human proteins as Zika antibodies, but through different pentamer motifs. For the envelope protein, these are shown in Table 2. Of particular interest are the human proteins where ZIKV has a high probability mimic. Most lineages of dengue type 3 carry a predicted mimic pentamer, GEDAP, which matches a pentamer in the mature NPY peptide. This differs from ZIKV, where the conserved pentamer mimic, ESTEN, is in the CPON component of the NPY propeptide [39], as shown in Figure 6. The dengue type 3 GEDAP motif is present in 134 of 277 recent (since 2000) South American dengue type 3 isolates analyzed. These subtype III isolates are from the Brazilian lineage. The remaining isolates, from the Venezuelan sublineage [40] instead carry the motif GEDVP. GEDVP has no mimic matches with proteins of neurologic function. GEDAP is conserved in all of 245 Asian dengue type 3 examined from the same time period. The proNPY mimic peptide in ZIKV is in EDIII and is strongly associated with predicted MHC II binding. In dengue type 3 GEDAP is located in EDIII at index position 322, a less exposed position, and adjacent peptides only have strong predicted binding affinity to 5 MHC II alleles. This implies there may be individual differences in antibody titer to this dengue epitope.

**Figure 6.**
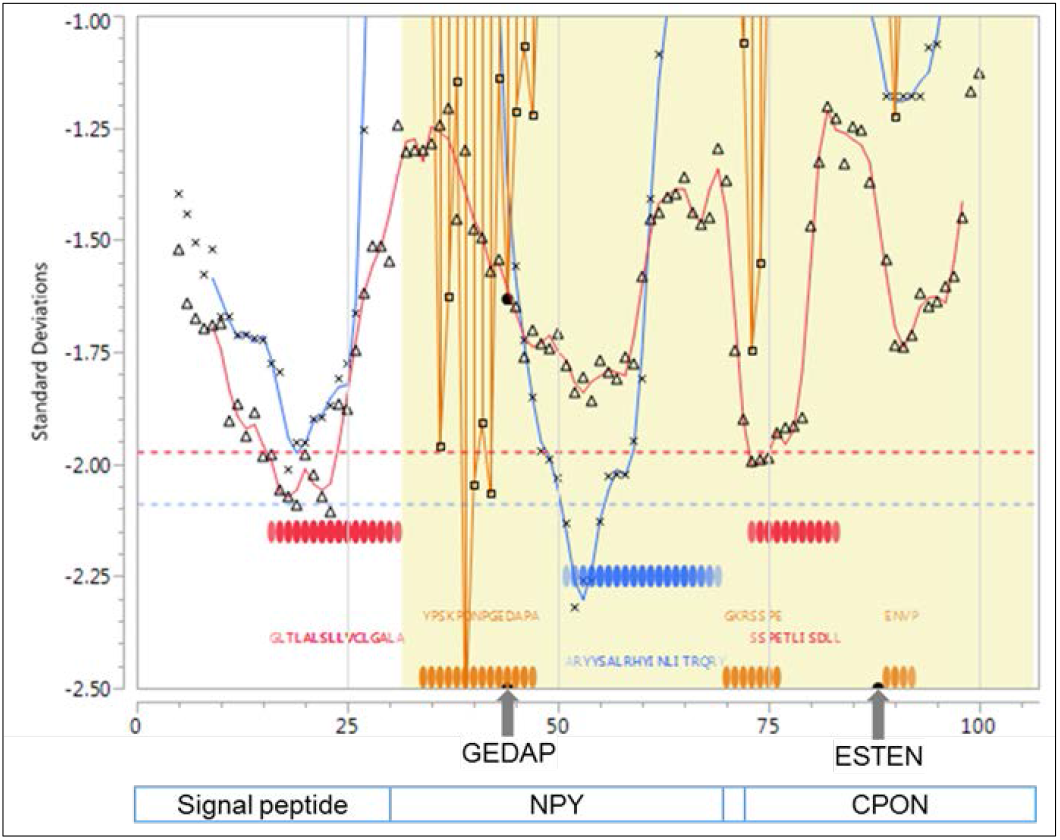
Immunome of pre-propeptide Y showing comparative positions of dengue 3 and Zika mimic pentamers. X axis shows N>C amino acid positions. Y axis shows standard deviation units of predicted MHC binding. Background shading shows signal peptide (white) and propeptide (yellow). Predicted MHC-I (red line), MHC-II (blue line) binding, and probability of B cell binding (orange lines) for each peptide, arrayed N-C, for a permuted population comprising 63 HLAs. Ribbons (red=MHC-I, blue-MHC-II) indicate the top 25% affinity binding. Orange bars indicate high probability B-cell binding. Arrows show predicted binding sites of antibodies from dengue 3 (GEDAP) and Zika (ESTEN).

**Table 2.**
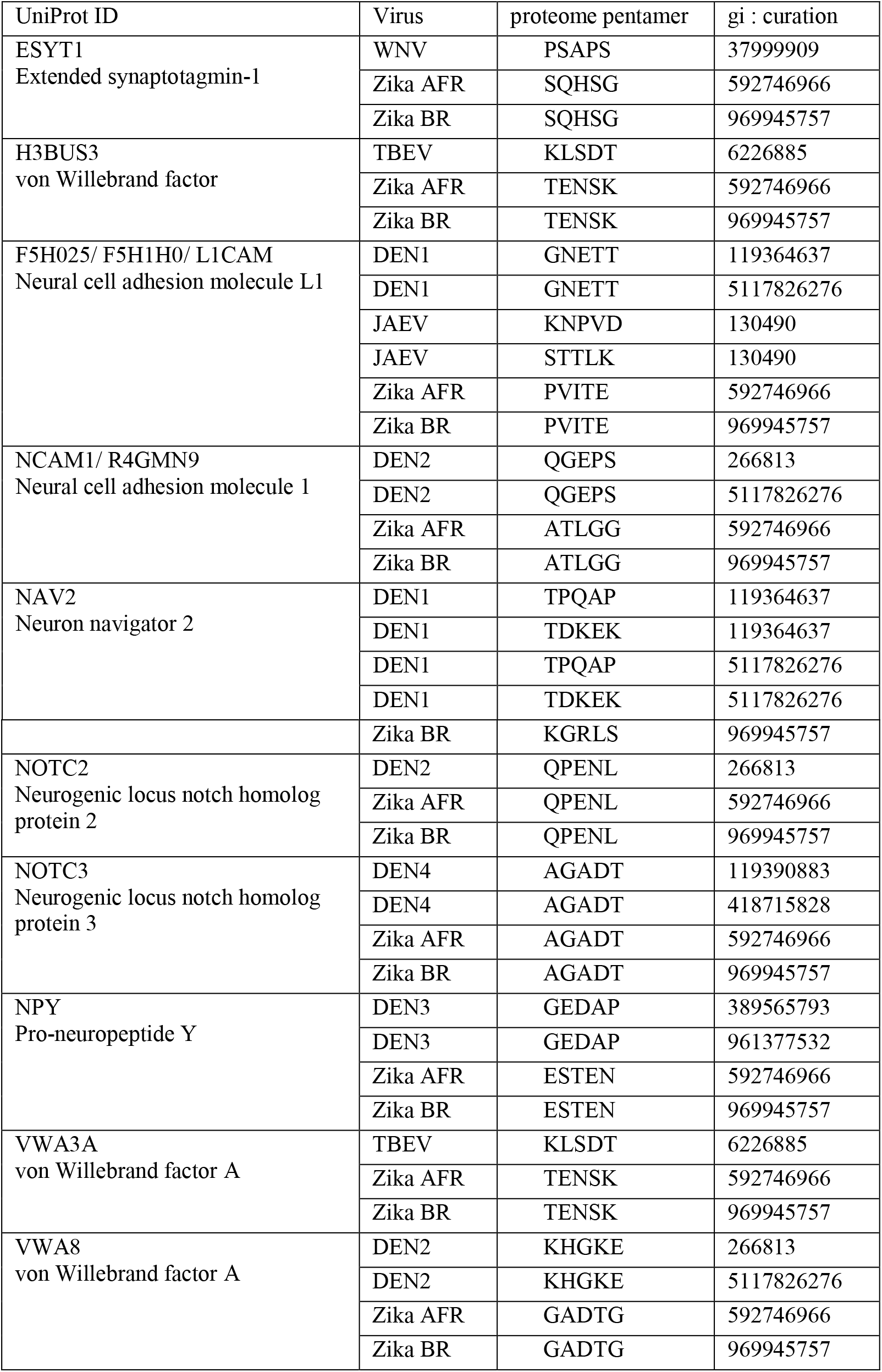
Proteins of neurologic function that are the targets of potential antibody mimicry by elicited by the envelopes of both Zika and other flaviviruses

Dengue type 1 carries two potential mimic pentamers for different locations in NAV2 (Figure 7). TPQAP is a low probability mimic due to its low BEPI score. TDKEK is in dengue type 1 EDIII DE loop and is present in all of 146 South American and 1126 Asian dengue type 1 isolates checked for the period 2000-2015.

**Figure 7.**
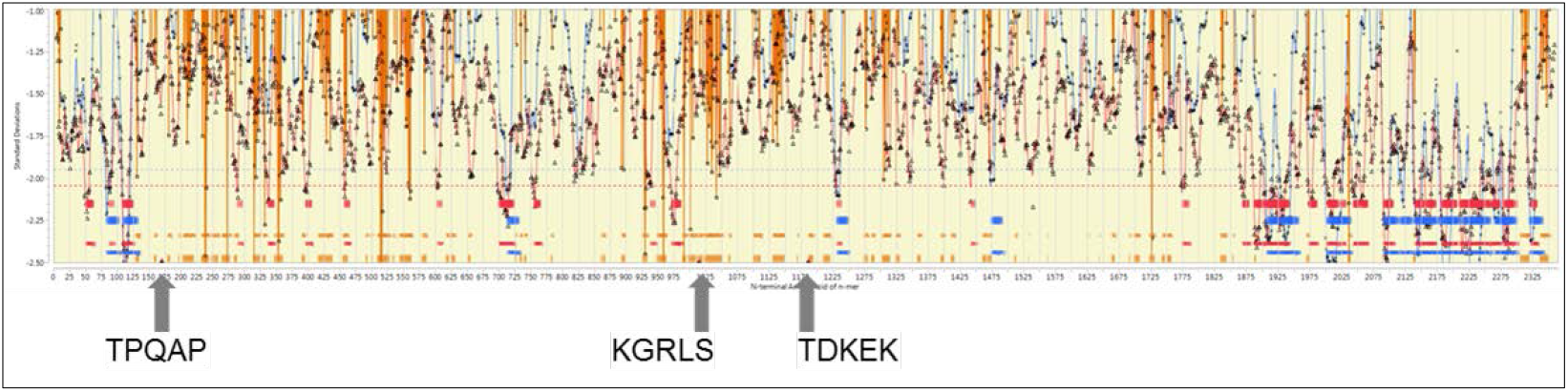
Immunome of NAV2 showing comparative positions of dengue 1 and Zika mimic pentamers. Axes and coloration as for Figures 1 and 6. Arrows show predicted binding sites of antibodies from dengue 1 (TPQAP and TDKEK) and Zika Brazil (KGRLS).

Our analysis of dengue viruses was done with the intention of identifying mimicry similarities and differences between dengue and ZIKV and included only a limited number of comparative strains of dengue virus. We did not conduct an analysis to identify all potential relevant mimics in dengue. In light of our findings, more in-depth analysis of these and other epitope mimics and their potential role in GBS as a sequel to dengue is warranted. Nevertheless, the fact that dengue type 3 and dengue type 1 also carry mimics on NPY and NAV2 respectively, suggests potential for an additive or complementary role to that of ZIKV. Other predicted mimics for other human proteins shared between ZIKV and other flaviviruses may have a combined impact. In each case this is conditioned by BEPI probability, and by whether associated T cell help enhances antibody titer.

### Other viruses

Preliminary analysis of chikungunya did not find potential neurologic protein mimics meeting our criteria in proteins E1 or E2, and one very low probability NAV2 match in the capsid, leading us to conclude that a cross-stimulation of Zika specific antibodies by this virus was unlikely. Comparative analysis performed on CMV and rubella was confined to surface and membrane proteins. This identified NAV2 matches in CMV membrane protein B and rubella E2. No NPY matches were detected.

## Discussion

Immunoinformatic analysis, such as that described here, has the advantage of rapidly bringing together a large array of data and enabling pattern recognition, often unapparent at the bench or in the field. It allows the development of hypotheses which must then be tested *in vivo*. In light of the severity of the threat posed by ZIKV, and the speed at which the epidemic is apparently expanding, there is an urgent need to develop medical countermeasures [41]. We therefore opted to make these results available to the Zika community to allow them to be considered before completion of field validation. This is currently limited by the biological time inherent in production of stringent test reagents.

Our analysis leads to the hypothesis that antibodies to ZIKV may contribute to the pathogenesis of the clinical outcomes, and indeed may be a primary driver of GBS, microcephaly, and the ocular lesions seen in infants. This has critical implications for development of therapeutic and prophylactic interventions. An antibody mediated effect may be independent of, or compound, the impact of viral replication.

Generally, the occurrence of potential mimicry, as laid out here for ZIKV, is no different from any other virus; all viruses carry pentamers which generate antibodies that react to varying degrees with targets on the human proteome. Some bind to target proteins having a critical function in a critical time interval; most do not. Secondly, BEPIs found in human proteins vary somewhat between isoforms, creating a range of probabilities (Supporting Figure S1). Yet another variable is the degree to which T cell help supports the generation of high antibody titers.

Thus, rather than a black and white/mimic versus not mimic, we are evaluating fuzzy logic, driven by overlapping probabilities. What sets ZIKV apart is that it appears to have a particularly problematic set of possible mimics. This includes the convergence of highly conserved potential mimics in its EDIII PVITESTENSK sequence, where there is contextually adjacent robust T cell help predicted. Added to this is the pair of newer high probability motifs in EDI which appeared with the F285S mutation.

Neutralizing antibodies for dengue typically cross link domains or molecules and limit movements of envelope proteins [34]. However, as evidenced by epitope mapping with peptides, this does not preclude linear epitopes from generating antibodies, neutralizing or not [36]. We would expect the same to be true for ZIKV. Coupling the amino acid sequence combinatorial selection with stringent co-selection of high BEPI probability, plus physiological and clinical criteria provides a very stringent and focused molecular identification process. This identifies a relatively small set of potential targets with likely clinical relevance. Our structural analysis of the consequences of the acquired mutations in the envelope identifies that the F285S mutation changes the exposure of BEPIs, possibly also creating new conformational epitopes. The impact of these mutations on the epitopes exposed in the immature virion, or their effect on constraining envelope flexibility, have not yet been explored.

The proteins identified as the targets of antibodies to high probability ZIKV mimic epitopes, including proNPY, NAV2, NDF4, NT4, BDNF, and neurexins, are proteins with diverse roles in neurologic function and in embryonic development[39, 42-46]. Sorting out the possible relative disruption of these in a ZIKV infection will be a complex challenge. We will only comment briefly on those proteins corresponding to the highest probability mimics.

NPY has been widely studied and found in many CNS and peripheral nervous tissue locations[47]. It is present very early in embryonic development[48]. Roles for NPY have been described in maintaining retinal development and health[39]. NPY is recognized as an immunomodulator with interactions with inflammation and immune response[49, 50] and is protective against the neurovirulence of retroviral infection[51]. The role of the CPON propeptide partner, where we predict anti-Zika antibodies would bind, is poorly understood, but CPON is widely distributed and contributes to neural injury recovery[52, 53]. Whether antibody binding would compromise propeptide processing is unknown, but is feasible. In an analogous situation, antibodies to the propeptide of insulin are important contributors to diabetes [54]. NPY is found in only one isoform in humans which, like insulin, may exacerbate the consequences of reduced NPY availability. Interestingly, the ESTEN motif is present in proNPY of primates, but variants occur in other species which may thus escape the effect of ZIKV mimics.

Optineurin, NAV2, and retinoic acid are part of the interactome necessary to retinal health and fetal CNS development [55-57]. In particular, NAV2 has been linked to fetal neurite growth and axonal extension [57]. Neural migration and extension are at a peak in the 13-21 weeks of pregnancy [58].

Antibodies binding to these proteins may compromise function or processing. In GBS, and other sensory deficits, the effect may be a transient impairment of neurologic function, corresponding to peak antibody titers. In a recent Zika report, a time line showed sensory impairment paralleling the antibody titer[3]. Interruption of function of one or more of the proteins identified during a critical stage of fetal development could lead to microcephaly and to the optic lesions observed following maternal ZIKV infection in pregnancy. This may be the result of placental transfer of maternal antibody, or endogenous fetal antibody, if fetal immune competence has been attained. An autoimmune response driven by antibody is not mutually exclusive with pathology arising from viral replication; both may occur.

Placental transfer of immunoglobulins to a fetus prior to blood brain barrier formation can be detrimental to the fetus. The human placenta facilitates the transfer of IgG, but not IgM, mediated by FcRn and increasing during the second trimester [59]. IgG1 and IgG4 are most efficiently transferred. Approximately 10% of maternal IgG is thought to pass into the fetal circulation, starting as early as week 13 [60]. The fetal blood brain barrier is not fully developed until the third trimester and may preferentially transfer proteins to the fetal brain [61, 62]. Thus, the literature suggests that the developing CNS is exposed to maternal antibodies in the first two trimesters. There is precedent for autoimmune diseases caused by the transplacental passage of antibody, including pemphigus, myasthenia gravis, and lupus [61, 63]. In dengue infection, maternal antibodies transfer to the fetus, achieving a level determined by maternal antibody titer [64]. Fetal anti-dengue titer may exceed maternal titer suggesting an active transfer process, without direct adverse effects on the fetus being reported until ADE following post-natal dengue infection [65]. A factor particularly enhancing placental transfer in the case of anti-Zika antibodies to the pentamers in PVITESTENSK may be enhanced immunoglobulin class switching to IgG, due to the adjacent MHCII binding peptides and T helper stimulation.

One of the vexing questions in Zika epidemiology has been why the observed pathology in the New World differs from that in Africa. While ZIKV is confronting a new, immunologically naïve population, the virus has also changed. As we show here the potential “mimicome” of ZIKV has evolved with each mutation. While some pentamers, such as those in the PVITESTENSK set, are conserved across all Zika isolates, others were more recently acquired. Microcephaly has not been reported in Africa, but was reported following the Zika outbreak in French Polynesia [66]. This follows the F285S mutation and associated predicted envelope mimic addition. Whether GBS has been associated with ZIKV over the years is unclear, as it is less conspicuous than microcephaly in an endemic setting, and especially in regions also endemic for trypanosomiasis, malaria, and dengue. This begs the question of whether this is a reporting phenomenon or whether some potential mimics may be more associated with GBS (e.g. conserved NPY matches) than microcephaly. Unravelling such complex interactions will take time.

The observation of high probability BEPIs, predicted to elicit antibodies to even higher probability BEPIs in VWA is an interesting coincidental finding. The relevant pentamer motifs are conserved between Zika Africa and Zika Brazil. The rash associated with ZIKV occurs after peak viremia, coincident with the initial rise in antibodies. This association may be multifactorial; antibody mimicry with other clotting factors has been noted in dengue [67], as well as depletion of VWA [68]. It is unknown what the impact of micro-hemorrhagic lesions in the fetus might be, given the greater vessel fragility [62].

Prior exposure to dengue can result in long duration of detectable antibodies [33]. All four serotypes of dengue circulate in the Latin American region currently affected by Zika virus, fluctuating both temporally and geographically. Co-circulation of multiple lineages of the same dengue type has been noted in Colombia [69]. The 2013 outbreak of Zika infection in French Polynesia was immediately preceded by reintroduction to the islands of dengue type 1 of SE Asian origin, and dengue type 3 subtype III of South American origin, but unspecified sublineage [2, 70]. Based on our findings, interactions with dengue could occur in two ways. As generally predicted, ADE arising from the conserved region of the envelope protein and PrM could exacerbate ZIKV infection. However, a further level of interaction may occur by the “doubling up” of mimics, as exemplified by NPY and NAV2, at different locations in each, by dengue 1 and dengue 3. Although there is overlap in the proteins targeted, the fact that anti-Zika antibodies arise from an epitope having strong predicted T helper support, may lead to higher titers and class switch than the antibodies from dengue epitopes with less T cell help. In this preliminary analysis we have only begun to address the potential interactions with dengue. Patterns of interaction with dengue may be particularly complex when multiple lineages co-circulate, some with and some without the NPY mimic [69]. A preliminary search of the envelope (E1, E2) and capsid proteins of chikungunya virus did not identify any strong potential mimics for the neurologically active proteins noted in ZIKV and dengue.

The pathology of microcephaly following Zika most closely resembles that observed in CMV and rubella infection [71]. While in both these cases virus has been isolated from the affected fetus and/or placenta, the exact pathway of pathogenesis is not understood [71, 72]. In CMV higher levels of fetal antibody were observed in infants born with sequelae of infection [73]. Our analysis of a small set of E1, E2, and capsid proteins of rubella, and the principal membrane proteins of CMV identified pentamer matches to NAV2, a finding that begs further inquiry.

While a causal Zika-GBS-microcephaly relationship appears increasingly, likely but awaits confirmation, the pathway laid out here provides a hypothesis for a mechanism of action and, consequently, how such a mechanism could be tested. The prospect that the pathogenesis of ZIKV may involve an antibody mediated autoimmune component must be addressed in vaccine and therapeutic development. In assessing the consequences of both vaccination and natural infection, it will be of particular importance to determine the duration of epitope-specific responses, as well as which epitope-specific responses are induced as an anamnestic response on re-exposure to the virus. Better understanding of epitope mimics may offer a specific pathway to GBS interventions, while clarification of the causal relationships of microcephaly in Zika infection may shed light on the pathogenesis of other viral teratologies.

## Acknowledgements

No external funding sources supported this study.

The authors gratefully acknowledge the comments and review of Drs. Michael Imboden, Stefanie Hone, Guy Plunkett III, Gary Splitter, and Thomas Yuill, and discussions with Drs. Adriano Schneider and Daniel Janies. Dr. Jill Glasspool Malone and Alleen Hager assisted with data compilation and literature assembly.

## Supporting Information

Figure S1 Correlation between the B cell epitope predictions for pentamers in the proteome corresponding to the identical pentamer in the Zika virus

Table S1: Viruses analyzed

Table S2: Keywords

Table S3: Predicted epitope mimics in all proteins of Zika Brazil

